# A low-cost, open-source framework for tracking and behavioural analysis of animals in aquatic ecosystems

**DOI:** 10.1101/571232

**Authors:** Fritz A. Francisco, Paul Nührenberg, Alex L. Jordan

**Affiliations:** Department of Collective Behavior, Max Planck Institute for Ornithology, Am Obstberg 1, 78315, Radolfzell, Germany; Centre for the Advanced Study of Collective Behaviour, University of Konstanz, Universitatsstraße 10, 78457, Konstanz, Germany; Department of Biology, University of Konstanz, Universitatsstraße 10, 78457, Konstanz, Germany

## Abstract

Although methods for tracking animals underwater exist, they frequently involve costly infrastructure investment, or capture and manipulation of animals to affix or implant tags. These practical concerns limit the taxonomic coverage of aquatic movement ecology studies and implementation in areas where high infrastructure investment is impossible. Here we present a method based on deep-learning and structure-from-motion, with which we can accurately determine the 3D location of animals, the structure of the environment in which they are moving. Further behavioural decomposition of the body position and contour of animals subsequently allow quantifying the behavioural states of each interacting animal. This approach can be used with minimal infrastructure and without confining animals to to a fixed area, or capturing and interfering with them in any way. With this approach, we are able to track single individuals (Conger Eel, *Conger oceanus*), small heterospecific groups (*Mullus surmuletus, Diplodus sp*.), and schools of animals (Tanganyikan cichlids *Lamprologus callipterus*) in freshwater and marine systems, and in habitats ranging in environmental complexity. Positional information was highly accurate, with errors as low as 1.67% of body length. Tracking data was embedded in 3D environmental models that could be used to examine collective decision making, obstacle avoidance, and visual connectivity of groups. By analyzing body contour and position, we were also able to use unsupervised classification to quantify the kinematic behavioural states of each animal. The proposed framework allows us to understand animal behaviour in aquatic systems at an unprecedented resolution and a fraction of the cost of established methodologies, with minimal domain expertise at the data acquisition or analysis phase required. Implementing this method, research can be conducted in a wide range of field contexts to collect laboratory standard data, vastly expanding both the taxonomic and environmental coverage of quantitative animal movement analysis with a low-cost, open-source solution.

## Introduction

Understanding the movement and behaviour of animals in their natural habitats is the ultimate goal of behavioural and movement ecology. By situating our studies in the natural world, we have the potential to uncover the natural processes of selection acting on the behaviour in natural populations, in a manner that cannot be achieved through lab studies alone. The ongoing advance of animal tracking and biologging has the potential to revolutionize not only the scale of data collected from wild systems, but also the types of questions that can subsequently be answered. Incorporating geographical data has already given insights, for example, into the homing behaviour of reef fish, migratory patterns of birds, or the breeding site specificity of sea turtles [7, 17, 46]. Great advances in systems biology have further been made through the study of movement ecology, understanding migratory patterns of birds traversing their physical environment or the decision-making processes at play within primate groups maneuvering through difficult terrain [36, 48]. Understanding these aspects of animal movement can also vastly improve management strategies [8, 9], for example in the creation of protected areas that incorporate bird migratory routes [43] or by reducing by-catch with dynamic habitat usage models [31].

Yet the application of techniques that meet the challenges of working in naturally complex environments is not straightforward, with practical, financial, and analytical issues often precluding effective usage and uptake of these approaches. This problem is disproportionately represented in certain ecosystems, with accessible solutions existing for some that simply do not work in others - for example the Global Positioning System (GPS) being very effective over savanna but failing entirely in underwater applications. These technical limitations can ultimately affect our understanding of entire ecosystems, with knock-on effects to all areas of knowledge and management. This becomes a fundamental problem if certain ecosystems, species, or habitat types fall behind the advances possible in other systems, because as the information available becomes limiting so too do options for informed management and discovery.

The inequality of tracking and animal movement approaches is perhaps nowhere better represented than by our lack of understanding of the oceans. Although the oceans constitute up to 90% of habitable ecosystems worldwide, as little as 5% have been explored [20, 34, 37]. Within the oceans, coastal inshore areas have the greatest species diversity, with approximately 80% of fish species (the most speciose group of vertebrates) inhabiting the shallow, littoral zone [41], while providing over 75% of commercial seafood landings [15]. Coastal regions in both marine and freshwater environments are also those that are of greatest interest for eco-tourism, community fisheries, and industry, while simultaneously being most affected by habitat degradation, exploitation, and anthropogenic pollution.

Yet aquatic ecosystems appear to be poorly represented in movement ecology research with only nine publications in the journal Movement Ecology containing one of the following keywords in the title: ‘ocean’, ‘aquatic’, ‘littoral’, ‘marine’, ‘fish’, ‘sea’[21]. Moreover, of the research into marine or freshwater animal movement, there is a heavy bias towards larger marine fauna, which often inhabit open oceanic areas [21]. This gap of knowledge comes as a surprise when considering the vast attention coastal marine systems receive in other areas of economic and usage management. Knowledge of the coastal regions is essential for establishing sanctuaries and sustainable concepts of ocean preservation [16] and movement data plays a vital role in this process, in that it gives detailed information about the location, preferred habitat and temporal distribution of organisms [28].

In the context of animal tracking, the tendency for studies to focus on larger and more charismatic animals is understandable from both a technical and engagement perspective. The noticeable size bias is mostly caused by the technical constraints of applied tracking methods, many of which traditionally rely on the application of tags that generate positional data that is most informative for larger animals. Tracking of tagged animals via GPS allows a sparse positioning of animals that surface more or less frequently, while Pop-up satellite archival tags (PSATs) integrate surface positions with logged gyroscope and accelerometer data for underwater position estimates [49]. Not only does the spatial resolution of respective tracking systems, e.g. 4.9m for GPS, limit the possibilities of behavioral analyses on a fine scale, but also excludes densely interacting animals from these research approaches [51]. Alternatively, ultrasonic acoustic telemetry can be used for underwater tracking of smaller animals and those in larger groups [30]. However, this approach is limited to stationary sites for positioning of the acoustic receivers. Further, the cost, maintenance, and installation of these systems preclude their effective use in the majority of coastal systems and for most users. Additionally, these methods require animals to be captured and equipped with tags that should not exceed 5% of the animals weight [10, 28, 30], rendering current GPS and PSAT tags problematic for small animals. While acoustic tags are small enough for injection, the increased handling time associated with these invasive measures can lead to additional stress for the animals, while the tag itself may disturb the animals’ natural behaviour. Hence, approaches that facilitate the collection of behavioural data in smaller animals, which compose the bulk of all species, are required.

The second source of this bias towards larger species is potentially associated with the greater engagement of the public funding bodies with charismatic megafauna. While we do not make any judgments about the value of studying one taxon over another, the benefits of techniques that are applicable to more species are beneficial in opening up avenues of novel research and insight, which may both contribute to a greater public understanding of ecosystems, and help reveal unanticipated behavioural and cognitive abilities in taxa such as fish [6, 29, 38]. Moreover, many of the established model organisms are small, and the combination of approaches from highly quantitative laboratory analyses with studies of natural ecology would open synergistic avenues of research in species in which we currently lack information about their natural behaviour and movement [47].

Overall, the benefits of applying these tracking and behavioural analysis techniques in a flexible, accessible, and broadly applicable manner provide ample reward for working to overcome limitations in systems and species coverage. This will improve conservation, management, and scientific understanding of natural systems across scales and conditions. Comparison of patterns of movement and search behaviour across a wider range of taxa may also reveal common rules to shared ecological problems, opening the potential for a fundamental understanding of the mechanisms and evolutionary origins of movement [21, 25]. Finally, with the application of quantitative behaviour and movement analyses in natural settings, the trade-off between high-resolution but contrived lab data and lower-resolution but naturalistic field data is lessened. Recent advances in behavioural decomposition [3, 23] and network analysis [11, 12] may then be employed in field settings, vastly improving our understanding of behaviour and movement in the wild [36].

In this paper, we present an open-source, low-cost approach based on consumer grade cameras to understand the movement and behaviour of animals of almost any size, as well as reconstruct the traversed environment, in coastal marine and freshwater ecosystems. Our approach synthesizes existing methodologies in machine-vision, neural network based machine learning, behavioural decomposition, and network analyses into a coherent framework that can be deployed in a variety of systems without domain-specific expertise. Object detection is achieved through computer vision [24], which has been successfully employed in terrestrial systems, through e.g. drone based video, yielding highly resolved movement data at the fine scale over broad environmental contexts. In addition to animal trajectories, this method also provides environmental data that adds the possibility to study interactions of mobile animals with their natural habitat [48]. While aerial drone-based approaches may also be used in some aquatic systems, they are limited to extremely shallow water and large animals [40] whereas the application we advocate allows data to be collected on any animal that can be visualized with cameras, hence potentially down to the millimetre scale. Subsequently, calculation of movement, interactions, and postures of animals, in combination with the construction of 3D models of the terrain with which animals interact is achieved through the open-source analysis pathway. Set-up costs can be as small as mobile phone cameras in waterproof bags, and can be taken into habitats which are otherwise explored by snorkeling, diving, or with the use of remotely operated underwater vehicles (ROVs). Analysis can be performed on open-source computing services or clusters (e.g. Google Colaboratory) or local GPU-accelerated machines. Overall, this method provides a low-cost approach for measuring the movement and behaviour of aquatic animals that can be implemented across scales and contexts.

## Materials and Methods

Three data sets of varying complexity were used to demonstrate the versatility of the proposed method. These were chosen to range from single animals (*'single'; Conger oceanicus*) and small heterospecific groups (*‘mixed’; Mullus surmuletus, Diplodus sp*.) to schools of conspecific individuals (*‘school’; Lamprologus callipterus*) under simple and complex environmental conditions, resulting in the data sets *single, mixed* and *school* respectively. The *single* and *mixed* data sets were created while snorkeling at the surface, using a stereo camera set-up (2x GoPro) at STARESO, Corsica (Submarine and Oceanographic Research Station). The *school* data set was collected via SCUBA (5-8 m) with a multi-camera array (12x GoPro) in Lake Tanganyika, Zambia (Tanganyika Science Lodge, Mpulungu). In contrast to the *single* and *mixed* data sets on untagged individuals, tags made of waterproof paper (8×8 mm) were attached anterior to the dorsal fin of the fish for the *school* data set [18]. Observations done prior suggest that these tags do not modify behaviour in comparison to untagged individuals.

### Automated animal detection and tracking

Since all data was collected in form of videos, animal tracking was required for subsequent behavioral analysis. Camera synchronization was achieved using a convolution of Fourier transformed audio signals to determine the video offsets in order to enable the advantages of a multiple view setup. The synchronized videos were tracked independently using an implementation of a Mask and Region based Convolution Neural Network (Mask R-CNN) for precise object detection at a temporal sampling rate of either 30 Hz (*single, mixed*) or 60 Hz (*school*) [1, 22]. Mask R-CNN models were trained on small subsets of labeled video frames with 40, 80 and 160 images for *single*, *mixed* and *school* respectively. The original image resolutions of 2704×1520 px (*single, school*) and 3840×2160 px (*mixed*) were downsampled during the training and prediction phase to achieve better performance. After training, the models were able to accurately detect and segment the observed animals, which was visually confirmed with predictions done on validation data sets.

The predicted masks were either used to estimate entire poses of the tracked animals (*single, mixed*) or to calculate centroids of the tags in case of the *school* data set. Established morphological image processing was used to skeletonize the binary mask predictions into approximations of the animals' spines, on which a fixed number of points was equidistantly distributed for fish pose estimation. Both the spine points and the tag centroid represent pixel coordinates of detected animals in further data processing. Partitioned trajectories were generated from detections with a simple combination of nearest-neighbor assignment and filtering for linear motion over a short time window, reducing later quality control and manual track identification for continuous trajectories to a minimum.

### Structure from motion

The field of computer vision has developed powerful techniques that have found applications in vastly different fields of science [14, 19, 52]. The concept of Structure-from-Motion (SfM) is one such method that addresses the large scale optimization problem of retrieving three dimensional information from planar images [32]. This approach relies on a static background scene, from which stationary features can be matched by observing them from different perspectives. This results in a set of images, in which the feature-rich key points are first detected and subsequently used to compute a 3D reconstruction of the scene and the corresponding view point positions. As shown in (1) and (2), a real world 3D point *M*′ (consisting of *x, y, z*) can be projected to the image plane of an observing camera by multiplying the camera’s intrinsic matrix *K* (consisting of focal lengths *f_x_*, *f_y_* and principal point *c_x_*, *c_y_*), with the camera’s joint rotation-translation matrix [*R*|*t*] and *M*′, resulting in the corresponding image point *m*′ (consisting of pixel coordinates *u, v*, scaled by *s*) [4]. By extension, this can be used to resolve the ray casting from a camera position towards the actual 3D coordinates of a point given the 2D image projection of that point with known camera parameters. Due to this projective geometry, it is not possible to infer at which depth a point is positioned on its ray from a single perspective. SfM is able to circumvent this problem by tracking mutually-observed image points (*m*′) across images of multiple camera view points. As a result, the points can be triangulated in 3D space (*M*′), representing the optimal intersections of their respective rays pointing from the cameras positions towards them. This approach is also able to numerically solve the multi-view system of the cameras relative rotation (*R*), translation (*t*) and intrinsic (*K*) matrices and to retrieve the optimal camera distortion parameters (*d*).

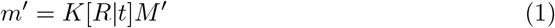

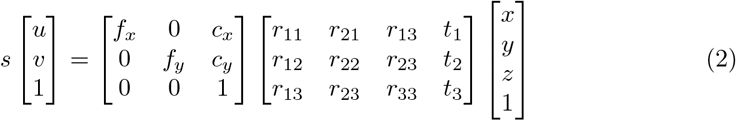

Here, SfM was incorporated into the process of data acquisition in order to gain information about the exact camera positions, which was done using the general-purpose and open-access pipeline COLMAP [44, 45]. The synchronized videos were split into images, serving as input for the reconstruction process, during which the cameras are calibrated (*K*, *d*) and relative extrinsic parameters (*R*, *t*) are computed, so that all camera projections relate to a shared coordinate system. Every image results in one corresponding point along the reconstructed, 3D camera path of the recording, where the number of images determines the temporal resolution of resolved camera motion. Only a subset of all video frames were used for reconstruction, sampling the videos with a rate of 3 Hz. This reduces the computational load, since COLMAP optimizes a smaller number of parameters. In addition, this could improve reconstruction accuracy, as the images still had sufficient visual overlap, but increased angles between viewpoints. The retrieved camera parameters were then interpolated to match the acquisition rate of animal tracking (60 Hz), assuring that reference camera parameters are given for each recorded data point by simulating a continuous camera path.

### Reconstruction of animal trajectories

It is necessary to resolve the camera motion in order to track moving animals with non-stationary cameras, since the camera motion will also be represented in the pixel coordinate trajectories of the animals. With camera information (*K*, *d*) and relative perspective transformations (*R*, *t*) for the entire camera paths retrieved from SfM, and multi-view animal trajectories available, a triangulation approach similar to SfM can be used to compute 3D animal trajectories. Positions of animals observed in exactly two cameras were triangulated using an OpenCV implementation of the direct linear transformation algorithm, while positions of animals observed in more than two cameras were triangulated using singular value decomposition following an OpenCV implementation [4, 19]. Additionally, positions of animals temporarily observed only in one camera were projected to the world coordinate frame by estimating the depth component as an interpolation of previous triangulation results. Through the recovered camera positions, the camera motion is nullified in the resulting 3D trajectories. Thus, they provide the same information as trajectories recorded with a fixed camera setup (Fig. 1). Animal trajectories and the corresponding reconstructions were scaled, so that the distances between the reconstructed camera locations equal the actual distances within the multi-view camera setup. As a result, all observations are represented on a real world scale.

**Figure 1.**
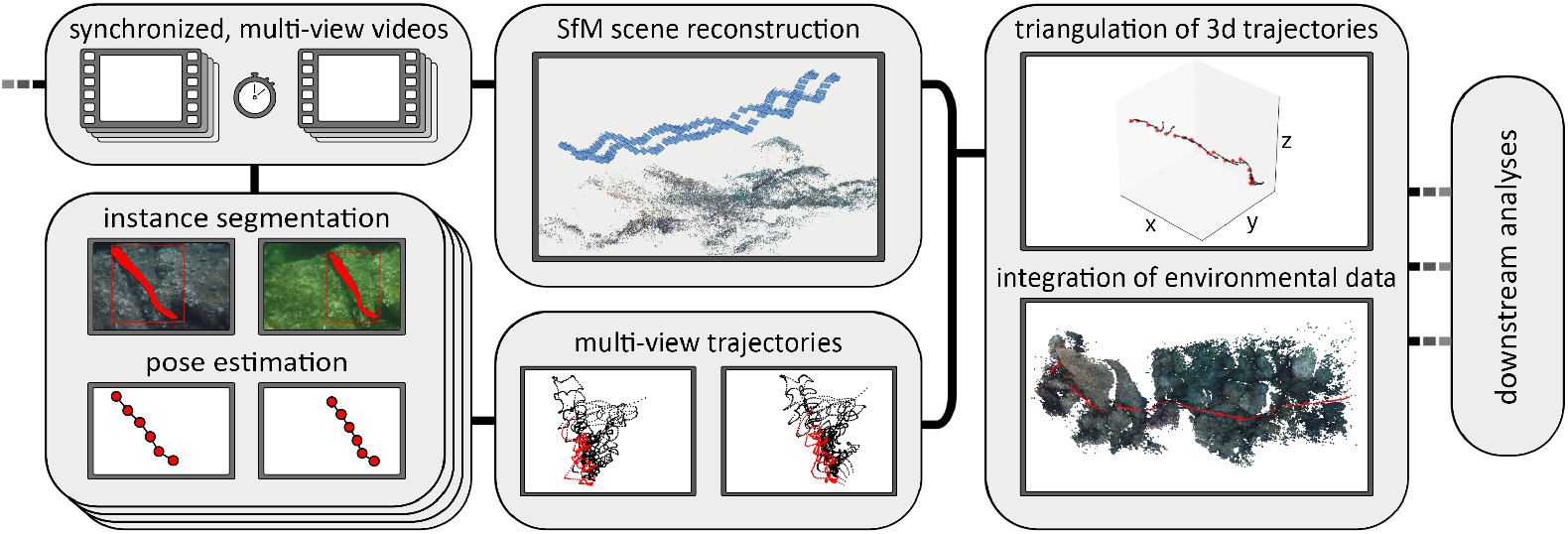
Schematic workflow. Data processing starts with acquisition of synchronized, multi-view videos, which serve as input to the SfM reconstruction pipeline to recover camera positions and movement. In addition, Mask R-CNN predictions trained on a subset of images result in segmented masks, from which animal poses can be estimated. They serve as locations of multi-view animals trajectories in the pixel coordinate system. These trajectories can be triangulated using the known camera parameters and positions from the SfM pipeline, yielding 3D animal trajectories and poses. Integrating the environmental information from the scene reconstruction, these data can be used for in depth downstream analyses.

## Results

Here we combine Mask-RCNN aided animal detection and tracking with SfM scene reconstruction and triangulation of 3D animal trajectories to obtain high resolution data directly from videos taken while snorkeling or diving in the field. Trajectories were successfully obtained from large groups *school*, small groups *mixed*, and single individuals *single*, and the environment through which animals were moving computed and reconstructed. Accurate estimation of fish body posture in 3D space for the *single*and *mixed* data sets (Fig. 3) was achieved by inference of spine points from the detection results (masks). Triangulation of the corresponding, multi-view animal detections resulted in small spatial errors (root-mean-square errors (RMSEs) 16.00 px, 5.27 px, 10.84 px of the trajectories 3D to 2D reprojections for the three data sets respectively *single, mixed, school*). This is equivalent to an average detection accuracy on the video images of 3.0% of the animals body length for the *mixed* and 1.67% length in the *single* data set. In the case of the *school* data set, which used tagging approaches, the average accuracy was lower at 29.4% of the tag diagonal (11.3 mm).

The acquired trajectories subsequently allowed to perform example quantitative downstream analyses developed in lab based applications. Trajectories themselves only represent one dimensional time series of the animals’ velocities, but estimated poses add a multitude of dimensions to these time series, such as the curvature or the width of the animals at every spine point. This data can be used for high throughput behavioral analyses based on unsupervised or supervised techniques [3, 26]. Here we used a simplified, unsupervised approach to classify motion states from trajectory data of the *school* data set. Wavelet transformations were applied on the time series of speed, acceleration and directional change to achieve a representation of animal motion in high-dimensional space. This was done in line with established methods, embedding instantaneous motion in broader time scales [3]. Non-linear dimensionality reduction was used to discretize this feature-rich data in a low dimensional embedding via FIt-SNE [33], resulting in motion states through which the animals cycle in a non-Markovian fashion. Animals were observed to show synchronization of these states, demonstrating that the data set could be used to infer the animals’ schooling properties (Fig. 2).

**Figure 2.**
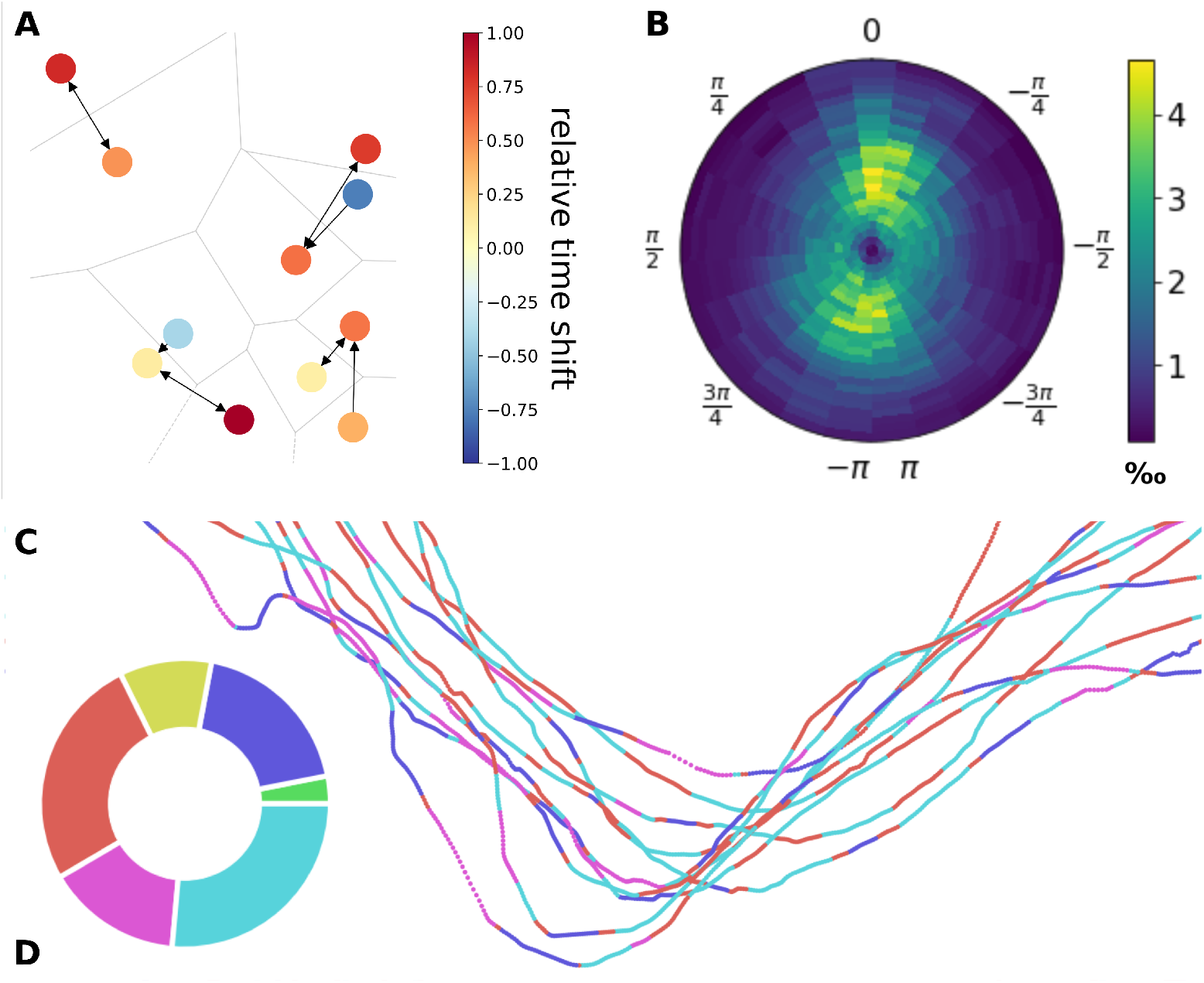
Downstream analyses. **A** Interaction network of the fish school at a single time step. Colors indicate the amount and sign of relative time shift to the maximally correlated individual (arrow). The Voronoi diagram gives an estimate of the group packing structure. **B** Polar probability density showing the likelihood of an individual to be present in any location (<0.5 m) relative to a focal individual. **C** Detailed view of trajectories, color coded by movement classification based on FIt-SNE embedding. **D** Overview showing the class proportions throughout the entire trial.

In order to resolve the interactions within the fish school, euclidean distances were calculated between individuals (*x, y, z* coordinates) resulting in a spatial proximity matrix for each frame. From this matrix, the substructure within the school was revealed through hierarchical clustering and by utilizing the elbow method to distinguish the optimal number of subgroups. Within these substructures the leader-follower relationships could be resolved over time through directional correlation analysis [35] (Fig. 2).

**Figure 3.**
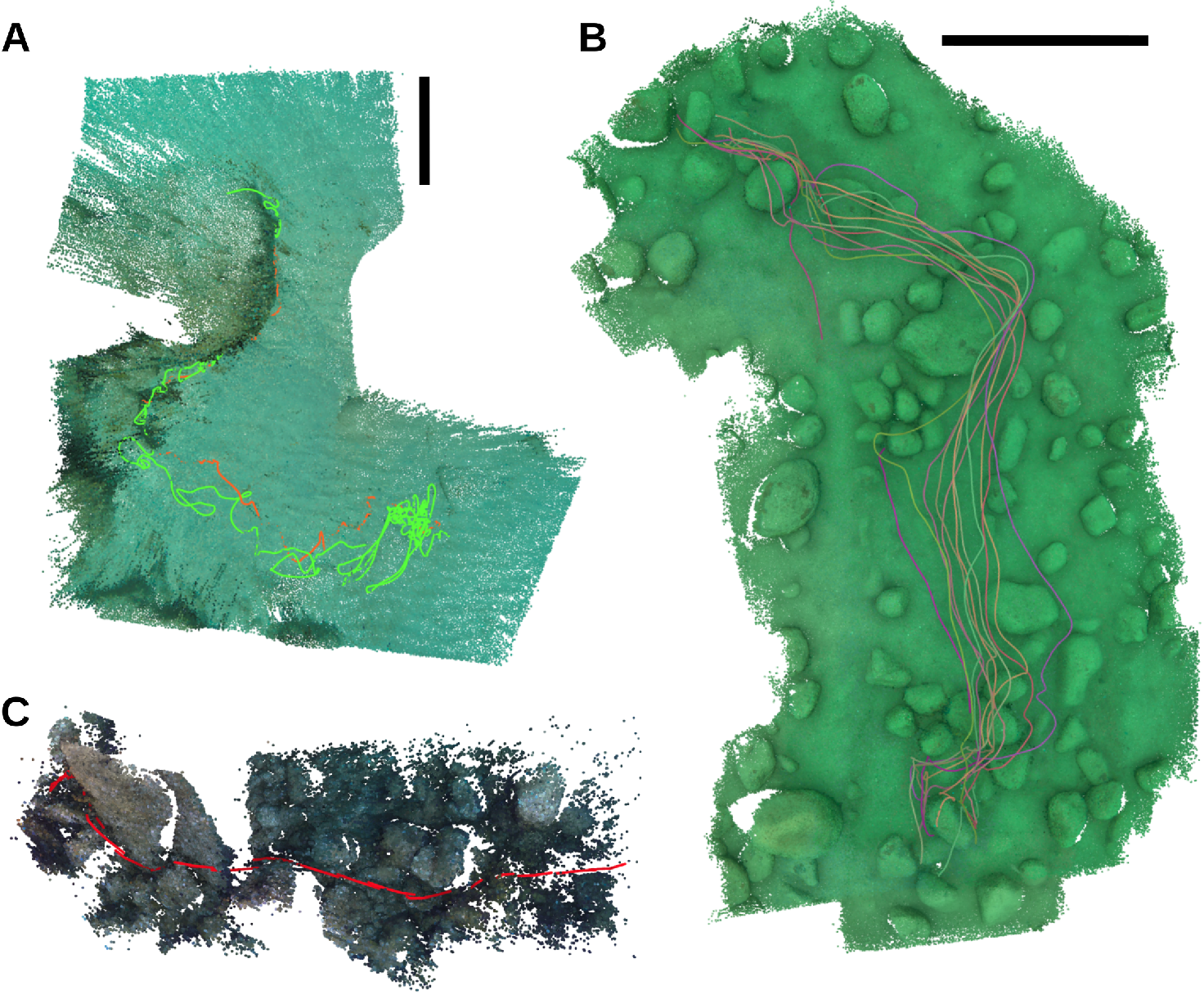
3D environments and animal trajectories. **A** Reconstructed environment as dense point cloud (green) resulting from high coverage footage, containing 12 individual tracks of *L. callipterus* (color) over a duration of 35 s. **B** Pose of single individual *C. oceanicus*. (red) shown every second moving through the environment reconstructed from sparsely populated coverage over a total of 20 s. **C** Trajectories of *M. surmuletus* (orange) and *Diplodus sp*. (green) over a duration of 7.1 min. Black bars denote scale (1 m). See additional Supporting Information for links to high resolution images.

## Discussion

The method we demonstrate here allows high resolution, 3D information of animal motion and interactions to be acquired in aquatic ecosystems. While the techniques are based on relatively advanced computational approaches, the open-source package we present requires little domain expertise and can be implemented with low-cost consumer grade cameras. The incorporation of these approaches will facilitate unprecedented applications for field based research across systems, scales, and hopefully users. Our approach allows data collection while swimming or snorkeling, and therefore makes it appropriate for general usage with minimal investment into infrastructure, equipment, or training. Although analyses are computationally demanding, they can be achieved on an average GPU or free cloud-based computing services. The lack of high-end hardware therefore does not interfere with any of the steps required for this method.

Although techniques for tracking of small aquatic animals do exist (e.g. telemetry, or in future, underwater time-of-flight cameras), these often have the considerable drawback of tagging and handling the animals or high infrastructure costs. In contrast, our approach does not require animals to be tagged, nor specialized equipment to be deployed. Our approach is also highly flexible to specific implementation requirements, for instance it can be used in clear water to resolve body posture and fine scale interactions, or can be combined with some form of tagging approach in conditions of high turbidity for example. Because the R-CNN approach can take any input, it is not tied to one particular animal shape or visual scene, and can therefore be flexibly used even in demanding conditions. While it is more limited in range, underwater filming comes as an unintrusive alternative to telemetry, and adds a data layer through the collection of environmental information. Although here we do not provide any analyses of environmental structure, this type of information is valuable when addressing questions on e.g. habitat segmentation and environmental complexity [13, 27].

While our approach offers many benefits in terms of applicability and data acquisition, it also suffers from some limitations. The SfM approach relies on the reconstructed components to be static, because key-points are assumed to have the same location over time. Any moving particles, besides the object of interest, will result in degradation of the reconstruction and higher reprojection errors. Very complex environments, occlusions of the animals and highly variable lighting conditions are detrimental to the detection ability and require consideration. The 3D pose estimation is highly reliant on accurate detections and can therefore be compromised by a poorly estimated animal shape during Mask R-CNN segmentation. In these cases, a less detailed approximation of the animals’ positions such as the mask centroid are favorable and can still be reliably employed as in the *school* example. The error in estimating animal location and pose can be partially explained by detection errors of the Mask R-CNN and inaccuracies derived from the trajectory triangulation. The latter can be estimated as RMSE of the reprojections from 3D back to 2D pixel coordinates. Substantial variation of error between body width and length (*single* data set) show that the reprojection error scales with object shape. Generally, error will increase with decreasing size of the object, although this size is only relative to the image frame itself, and so can be resolved with zoom lenses or close-up filming of smaller animals. In order to ground truth the 3D projection and resolve the total error, a calibrated, under-water space would be required. This was not possible in the presented trials since all data sets were recorded in the natural environment, allowing the animals to move freely without the boundaries of such a standardized space. Further, referencing the reconstructions to a metric scale is only applicable if the distance between cameras is known or the reconstructions contain objects of known size. This was only the case for two (*mixed, school*) of the three example data sets.

An additional limitation of our approach is associated with the need to annotate images and train detection networks. However, this additional time investment is subsequently offset by the time saved in subsequent observations by using automated detection and classification of behavioural states, for instance by quantifying the behavioural repertoire of the animal using unsupervised machine learning techniques [3, 42, 50]. The adaptation to three dimensional motion analyses have allowed for a better understanding of the phenotype and development of animal behaviours [53]. In addition, 3D pose estimation is now possible for wild animals, enabling exact reconstruction of the entire animal [54]. There has been a shift in how animal movement is analyzed in light of computational ethological approaches [5, 39, 42], with patterns of motion able to be objectively disentangled, revealing the underlying behavioural syntax to the observer. Automated approaches based on video, or even audio, recordings may also overcome sensory limitations of human systems, allowing a better understanding of the sensory *umwelt* of study species [25] and also facilitate novel experimental designs [2, 42] that can tackle questions of the proximate and ultimate causality of behaviour [5, 39, 54]. These methods are gaining interest and sharply contrast with the traditional approach of trained specialists creating behavioural ethograms, but can usefully be combined and compared to gain further insight into the structure of animal behaviour, potentially generating a more objective and standardized approach to the field of behavioural studies [5].

Advances in analytical prospects for behavioural studies in the lab, coupled with quantitative tracking approaches of animals in the wild, opens the possibility to advance our knowledge of natural systems with highly quantitative data streams [5, 28, 39]. However, for effective uptake of these techniques, a common, cost-efficient framework is required. Here, the acquisition of both animal movement and environmental information is combined into a single data collection pipeline. It is implemented in a cost-efficient and open-source fashion, yet it can be easily integrated into more exclusive deployment and remote sensing applications, such as deep sea ROVs, widening the possibilities in the studies of aquatic animals.

## Supporting Information

### 3D environment and track reconstructions

1. *single*: C. oceanicus @ STARESO, Corsica
2. *mixed*: M. surmuletus & Diplodus sp. @ STARESO, Corsica
3. *school*: L. callipterus @ Kasakalwe, Tanganyika

## Acknowledgments

We would like to thank the entire Department of Collective Behaviour at the University of Konstanz for their support in making this project possible. We thank Philip Fourmann, Myriam Knöpfle and Jessica Ruff for contributing the *single* and *mixed* species footage, and Maté Nagy for his help on the correlation analysis. Special thanks also to Hemal Naik, Simon Gingins and Eduardo Sampaio for their suggestions and helpful input. We sincerely thank Etienne Lein for his substantial assistance and support during data acquisition in the field. Further, we thank the COLMAP team for making it open-source and easily accessible.

